# Human primed endothelial colony forming cells exert neuroprotective effects in the growth restricted newborn piglet

**DOI:** 10.1101/2021.02.15.431307

**Authors:** Kirat K. Chand, Jatin Patel, Tracey Bjorkman, Seen-Ling Sim, Stephanie M. Miller, Elliot Teo, Lara Jones, Jane Sun, Paul B. Colditz, Kiarash Khosrotehrani, Julie A. Wixey

## Abstract

The fetal brain is particularly vulnerable to the detrimental effects of fetal growth restriction (FGR) with subsequent abnormal neurodevelopment being common. There are no current treatments to protect the FGR newborn from lifelong neurological disorders. This study examines whether pure fetal mesenchymal stem cells and endothelial colony forming cells (ECFC) from the human term placenta are neuroprotective through modulating neuroinflammation and supporting the brain vasculature. We determined that one dose of these primed ECFCs (pECFC) on the first day of life to the newborn FGR piglet improved damaged vasculature, restored the neurovascular unit, reduced brain inflammation and improved adverse neuronal and white matter changes present in the FGR newborn piglet brain. These findings could not be reproduced using mesenchymal stromal cells alone. These results demonstrate pECFC treatment exerts beneficial effects on multiple cellular components in the FGR brain and act as a neuroprotectant.

**One Sentence Summary:** Stem cell treatment improves brain outcomes in the growth restricted newborn

## Introduction

Fetal growth restriction (FGR) occurs in around 3-15% of pregnancies with even greater rates in developing countries (*1–3*). FGR is often caused by placental insufficiency, resulting in an inadequate supply of oxygen and nutrients to the developing fetus *in utero* (*4*). The fetal brain is particularly vulnerable to FGR conditions and subsequent abnormal neurodevelopment is common. Adverse neurodevelopmental outcomes include learning and behavioral disorders and cerebral palsy, which have lifelong medical and financial consequences (*5–9*). Due to medical advances, more FGR babies now survive, although many remain at risk of these neurodevelopmental disabilities. No treatments currently exist to protect the FGR newborn brain.

Both neuronal and white matter alterations are observed in both the human FGR infant and animal models of FGR (*10–15*). Recent animal studies have shown that inflammation plays a key role in these alterations (*16–18*). High concentrations of pro-inflammatory cytokines have been reported in the blood of preterm FGR infants two weeks after birth (*19*). Increases in pro-inflammatory cytokines are correlated with adverse neurological outcomes at 2 years of age in preterm small for gestational age newborns (*20*). An increase in inflammatory mediators in the blood may affect the brain because of the disruption to the structural and functional integrity of the neurovascular unit (NVU) in the FGR newborn (*21–23*) and toxic mediators may pass freely in and out of the brain. Therapeutic targeting of inflammatory pathways and the NVU hold promise as neuroprotectants in FGR newborns.

The NVU separates the brain from the blood circulation and is composed of vascular endothelial cells, glial cells (astrocytes and microglia), neurons, pericytes, and the basement membrane. The cells of the NVU share close and complex interactions that are crucial in maintaining blood-brain barrier (BBB) integrity and cerebral homeostasis, ensuring healthy brain development. Disruption to the NVU plays an essential role in progression of numerous central nervous system (CNS) pathologies allowing toxic substances, such as pro-inflammatory cytokines to infiltrate the brain, which can exacerbate neuroinflammation and injure neurons and white matter (*24, 25*). As the NVU is developed in the newborn, maintaining the structural and functional integrity of the NVU is likely an essential element in neuroprotection in the newborn.

Endothelial colony forming cells (ECFCs) are the vascular precursors, devoid of hematopoietic and myeloid contamination (*26*). They reside throughout the vasculature, have self-renewal as well as a capacity to engraft forming both *de novo* neo-vessels as well as chimeric vessels within ischemic sites of the host, facilitating reperfusion and initiating tissue regeneration (*27, 28*). They have been isolated from numerous sources, classically the umbilical cord blood (UCB) and the term placenta (*29, 30*). The vasculogenic capacity of ECFCs is greatly enhanced upon co-incubation or co-delivery with mesenchymal stromal cells (MSCs), a process known as priming (*31*). Moreover, engraftment of ECFCs if primed (delivered with MSCs) can bypass the host immune system (*32*). Recent evidence suggests that MSC/ECFC injected in combination, outperformed MSCs alone in reperfusing ischemic limbs through neo-vascularisation (*27*). Additionally, this combination of primed ECFCs did not require additional immunosuppressive therapy in immunocompetent animals (*27*) highlighting their potential use as an allogeneic off-the-shelf cell therapy.

In the present study, we obtained pure fetal MSC and ECFC from the healthy human term placenta (*30, 33*). We hypothesized that the primed ECFC (pECFC) treatment provides neuroprotection in the FGR newborn. Using a preclinical piglet model of FGR we have established (*16, 34, 35*), we assessed whether one dose on the first day of life in a term FGR piglet would regenerate damaged vasculature, restore the NVU, reduce brain inflammation and improve adverse neuronal and white matter changes present in the FGR newborn piglet brain.

## Results

Body weight and brain weight were significantly lower for all FGR piglet groups at postnatal day 4 (P4) compared with normally grown (NG) piglets (body p<0.0001; brain p<0.05; Table 1). Piglets in all FGR groups were asymmetric indicated by a mean brain to liver weight ratio (B:L) >1. There was no significant difference in body weight (p=0.735), brain weight (p=0.999) or liver weight (p=0.721) between NG piglet groups (Table 1).

**Table 1:** Physiological parameters in FGR and NG piglets. Piglet bodyweight, brain weight, liver weight, temperature, and mortality. Mean body weight was significantly lower in all FGR groups compared with NG piglets. Brain weight was significantly reduced in all FGR groups compared with the NG group. Mean brain to liver weight ratio was significantly higher in all FGR groups compared with NG piglets indicating asymmetric growth restriction in the FGR piglets. No significant difference in average temperature over the 3 days. Values are the mean±SEM. *p<0.05; **p<0.01; ****p<0.0001 versus NG.

|  | NG (n=8) | FGR (n=8) | FGR+pECFC (n=8) | NG+pECFC (n=7) | FGR+MSC (n=5) |
| --- | --- | --- | --- | --- | --- |
| Bodyweight in grams (mean±SEM) | 1968 ± 138.10 | 1023 ± 77.13**** | 1028 ± 88.96**** | 1743 ± 91.59 | 800 ± 35.78**** |
| Brain weight in grams (mean±SEM) | 32.74 ± 0.50 | 29.97 ± 0.60* | 29.38 ± 1.18* | 32.03 ± 0.59 | 28.87 ± 0.82* |
| Liver weight in grams (mean±SEM) | 51.24 ± 6.16 | 24.19 ± 1.86**** | 25.22 ± 2.77**** | 42.66 ± 2.94 | 21.49 ± 1.73**** |
| Brain:liver ratio (mean±SEM) | 0.69 ± 0.07 | 1.30 ± 0.12** | 1.29 ± 0.17** | 0.78 ± 0.06 | 1.38 ± 0.11** |
| Temperature (average over 3 days) | 38.33 ± 0.14 | 38.39 ± 0.08 | 38.15 ± 0.08 | 38.2 ± 0.09 | 38.03 ± 0.21 |
| Mortality | 0/8 | 1/9 | 0/8 | 0/7 | 0/5 |

### pECFC delivery and distribution

Placental ECFC and MSC were obtained from 3 donors. 20 piglets received intravenous (i.v.) stem cells (either pECFC or MSC) versus Sham. No differences in body temperature were evident between any of the five experimental groups (Table 1). Normal temperature for a piglet is approximately 38.5°C (*36*). Piglets in all five groups responded well to feeds with one death (due to inability to thrive) in the FGR group (Table 1).

Biodistribution of human stem cells were detected in the piglet brain at P4 based on labelling with a human specific Lamin A/C antibody. Cells positive for Lamin A/C were found to engraft into the vasculature of microvessels as well as sporadic instances of parenchymal labelling in FGR+pECFC brain (Figure 1C). No Lamin A/C labelling was observed in the technical control (no primary antibody) nor in the NG+pECFC piglet brain (Figure 1D & E). Overall, pECFC delivery was deemed safe and distributed to the vasculature and perivascular areas of the FGR brain.

**Figure 1.**
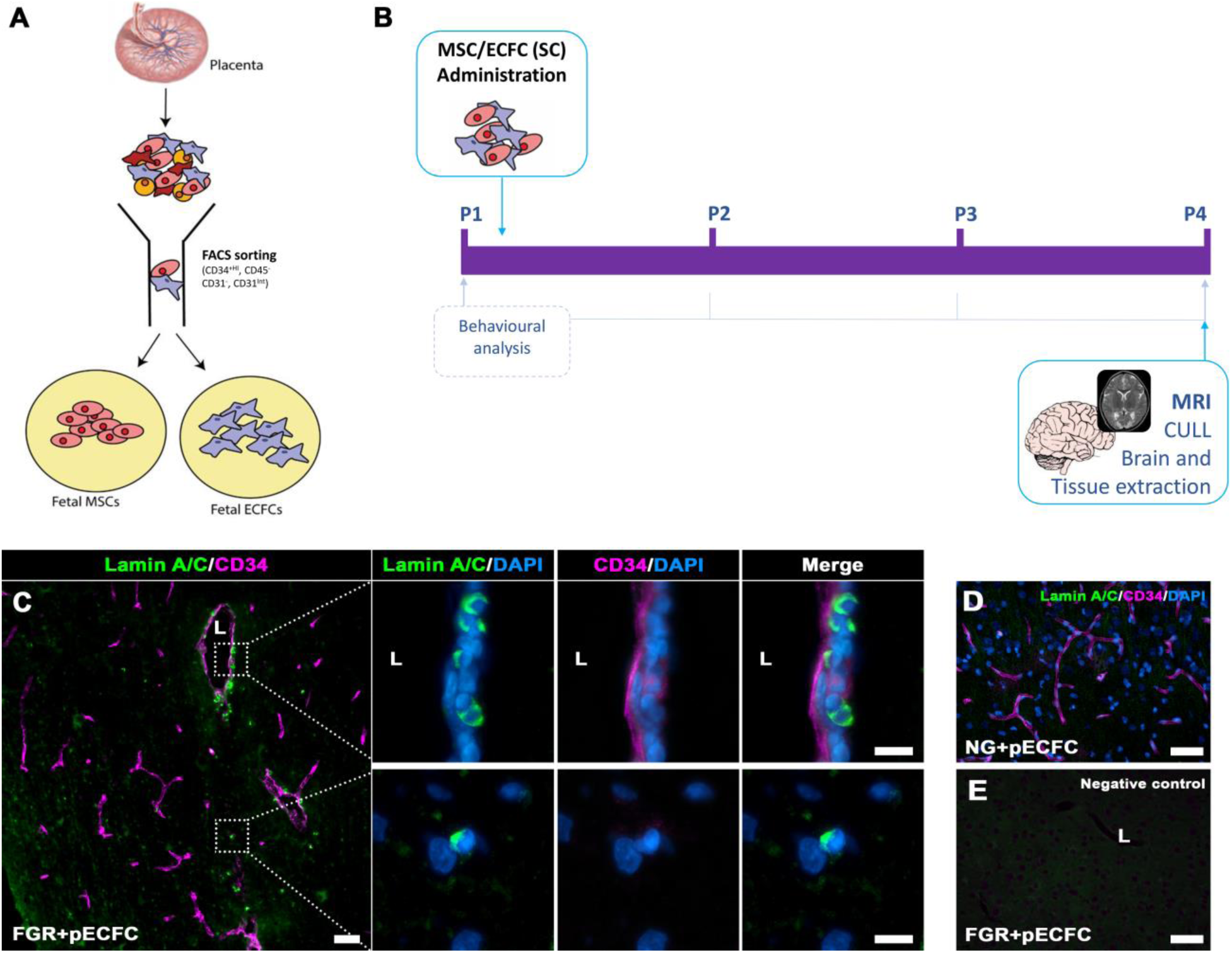
Isolation and administration of combined Mesenchymal and Endothelial Colony Forming Cells. **(*A*)** Fetal MSC and ECFCs were derived and isolated from healthy human placenta via FACs sorting. Isolated cells were cultured, enriched and prepared for intravenous administration as a combined SC preparation (10^6^ MSC and 10^6^ ECFC). **(*B*)** Schematic representation of MSC/ECFC therapy in the newborn FGR pig model. **(*C*)** Presence of cells in treated FGR brain was confirmed post-mortem (P4) using Lamin A/C. Lamin A/C-positive cells were observed in the parenchyma as well as instances of vessel engraftment (lumen; L). **(*D*)** No Lamin A/C-positive labelling was observed in NG+pECFC tissue and **(*E*)** negative control (primary antibody omitted) (Scale bars: 50µm; ***C*** high magnification: 10µm).

### pECFC administration promotes vessel density

We investigated the potential benefits of pECFCs in improving vasculature impairment of the FGR brain. A key component of the cerebrovasculature is collagen IV (Col IV), which contributes to approximately 50% of the basement membrane of the NVU. All groups displayed robust labelling of Col IV across the length of the vasculature, however areal density analysis showed a significant reduction in the FGR brain compared with NG (p=0.009; Figure 2A&B) indicating a reduction in vascularisation. pECFC-treated FGR brain displayed higher vessel density compared with FGR (p=0.048; Figure 2A&B). Administration of MSC had no significant effect on vessel density in FGR (Supplementary Figure 1A&B). Analysis of the endothelial marker (CD34) closely reflected the observations of Col IV labelling with significantly less labelling in FGR brains compared with NG (p=0.019; Figure 2C&D), that was normalised with pECFC treatment. Examination of vascular length and vascular complexity based on the number of branch points from the primary vessel revealed a significant decrease in the total vascular length in FGR brain compared with NG (p=0.025; Figure 2E) with significant restoration upon pECFC treatment (p=0.021; Figure 2E). In addition to loss in vessel length we also observed a decrease in vessel branch points in FGR brain relative to NG (p=0.002; Figure 2F). pECFC treatment partially improved vessel branching in FGR+pECFC brains appearing similar to that observed in NG brain (FGR+pECFC *c.f*. NG, p=0.405). Although we could not find any evidence of cells in the NG+pECFC piglet brain comparison of NG and NG+pECFC of vessel length and branching showed no differences suggesting pECFC administration does not promote unwarranted angiogenic effects. These findings support the potential of pECFC as a safe treatment to improve cerebrovascularisation in the FGR neonate as previously shown in other ischemic scenarios.

**Figure 2.**
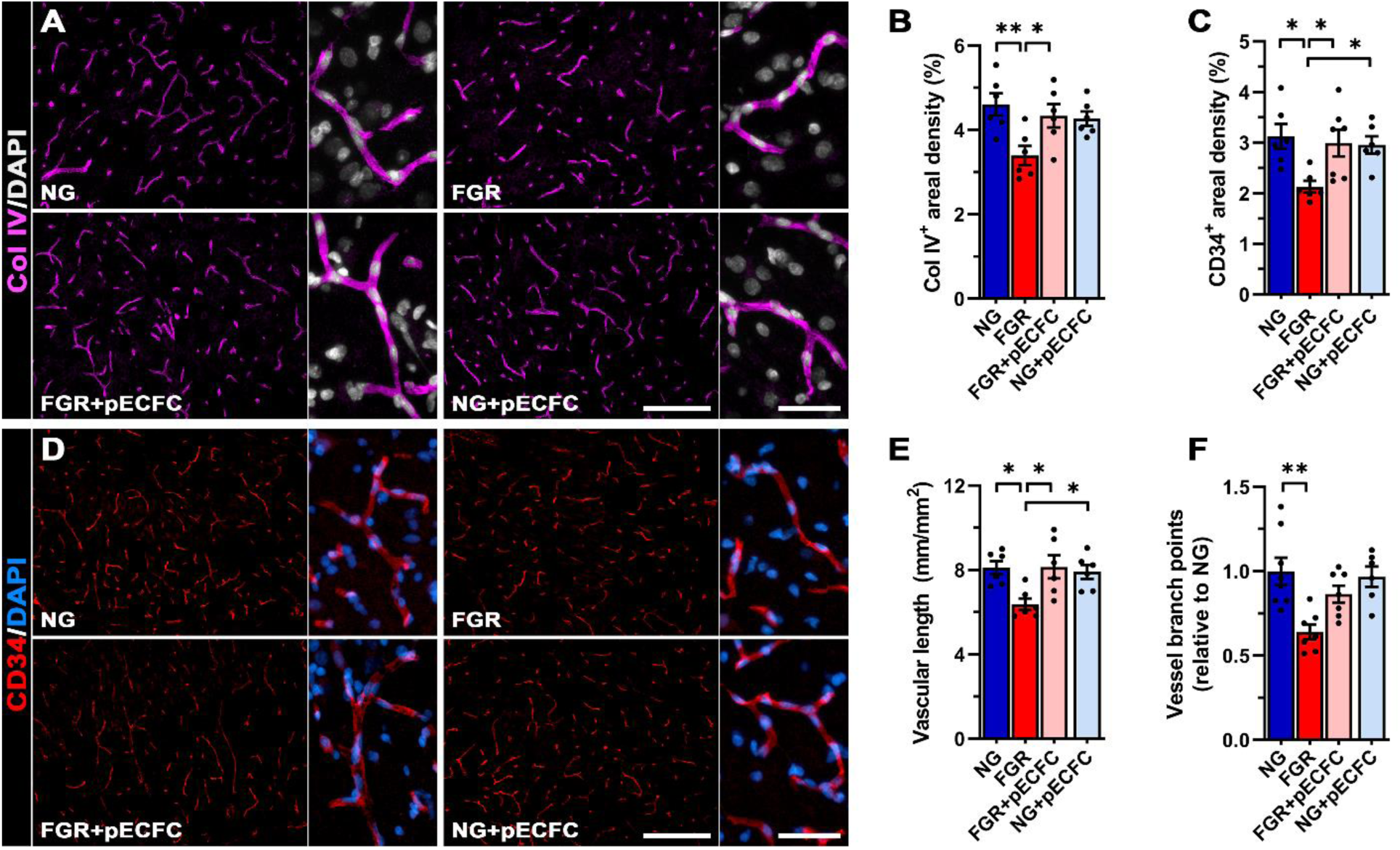
pECFC administration promotes vessel density and reduces BBB-permeability. **(*A*)** Representative immunofluorescent labelling of vessel basement membrane (Col IV; magenta) in the parietal cortex. FGR displayed truncated vasculature with limited branching compared with other groups. **(*B*)** Areal analysis found a significant decrease in coverage of Col IV-positive vessels which improved following pECFC administration. **(*D*)** Progenitor endothelial cell marker (CD34; red) showed similar patterns observed as Col IV, with FGR demonstrating lower areal coverage **(*C*)**. **(*E*)** Analysis of the vasculature showed a significant decrease in vessel length as assessed by labelling of Col IV. **(*F*)** Vessel branching was also reduced in FGR relative to NG. FGR+pECFC did not display increased vascular branching. All values are expressed as mean +/- SEM (minimum *n*=6 for all groups). Two-way *ANOVA* with Tukey post-hoc test (*p<0.05, **p<0.01) (Scale bars: low magnification: 200µm; high magnification: 50µm).

### pECFC treatment enhances neurovascular unit integrity

We further investigated whether the improved vascular structure and organisation in the FGR brain that occurred with pECFC treatment, resulted in maintenance of BBB integrity. Previous studies have demonstrated the utility of endogenous serum proteins as markers of altered BBB-permeability. We used albumin (∼69kDa) and IgG (∼155kDa) extravasation to assess whether pECFC-treated animals demonstrated improvement to BBB permeability. FGR brains displayed predominantly perivascular labelling of albumin (Figure 3A) with no observed extravasation into the parenchyma. Astrocyte end-feet encasing vessels with perivascular albumin presented activated morphology, with hypertrophic end-feet and swelling of cell bodies (Figure 3Aa-b’). In comparison, NG and pECFC-treated groups displayed low intensity albumin labelling that was predominantly restricted to the vessel lumen and a return of juxtavascular astrocytes to a ramified astrocyte morphology (Figure 3B, C & D). Quantification of albumin labelling showed a greater number of vessels with perivascular labelling in FGR compared with NG (p=0.007; Figure 3E). Albumin areal density, used to assess the aggregation of albumin in the perivascular space, was significantly higher in FGR when compared with NG (p<0.0001) and was significantly reduced upon pECFC treatment (p=0.009; Figure 3F). However despite pECFC treatment, albumin in the perivascular space remained significantly greater when compared with NG (p=0.005; Figure 3F).

**Figure 3.**
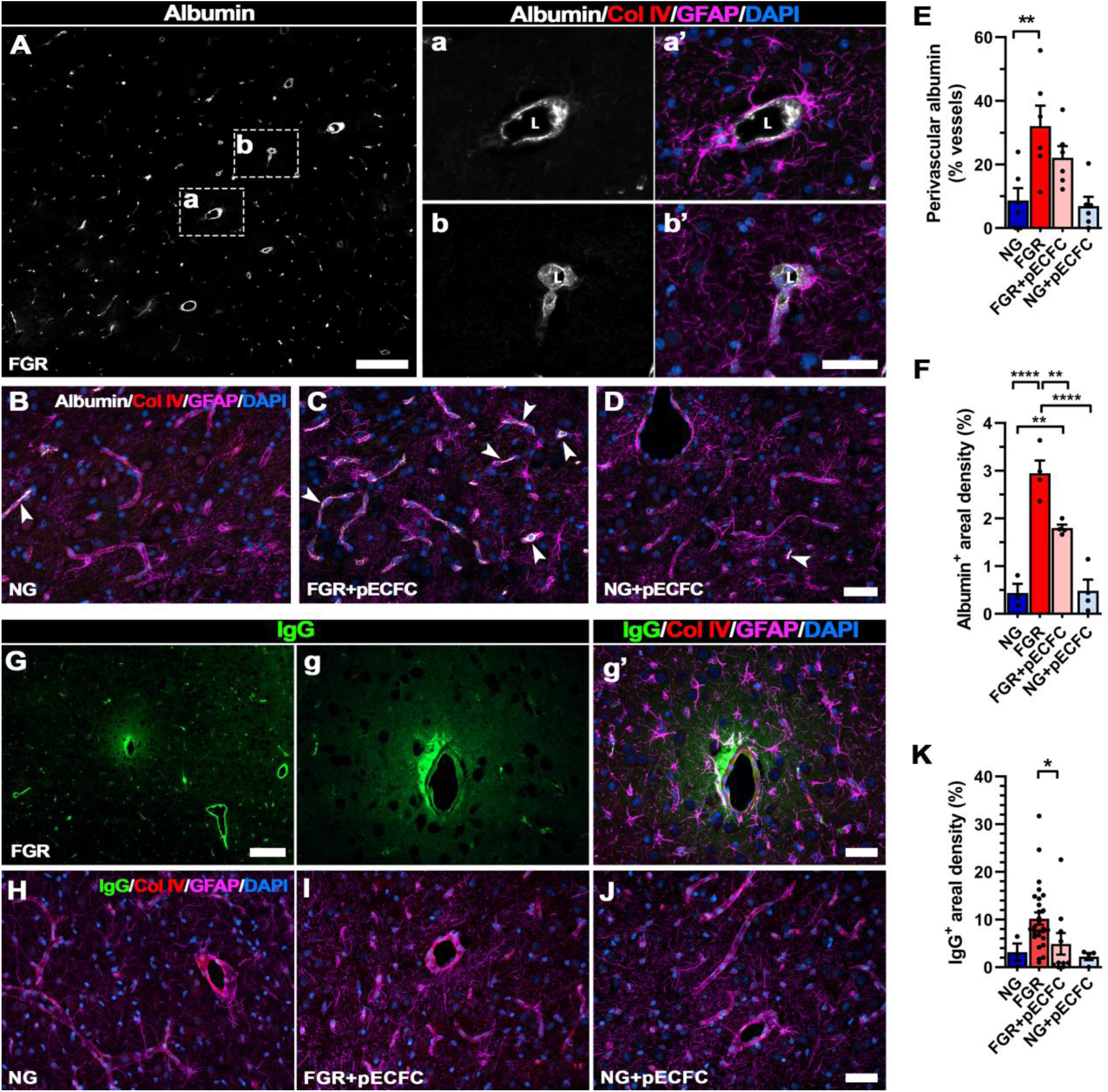
pECFC administration ameliorates neurovascular integrity in the FGR neonate. Representative labelling of endogenous albumin in the FGR as a marker of altered blood brain barrier integrity. **(*A*)** FGR brain showed albumin labelling (grey) predominantly localised to the perivascular space, between the lumen (L) and astrocyte endfeet (GFAP; magenta) (see ***Aa’-Ab’***). **(*B*)** NG, **(*C*)** FGR+pECFC and **(*D*)** NG+pECFC displayed lower less overt albumin labelling. **(*E*)** A significantly higher percentage of vessels in FGR brain displayed perivascular labelling compared with NG. Stem cell treatment did not significantly reduce the number of vessels with perivascular labelling. **(*F*)** FGR brain presented significantly greater areal density of albumin- positive labelling compared with NG. FGR+pECFC showed lower areal coverage but was still elevated compared with NG. **(*G*)** Representative labelling of IgG (green) in the FGR brain showed extravasation into the brain parenchyma. Evident astrocyte activation (GFAP; magenta) was observed a vessels displaying altered permeability. **(*H*)** NG and **(*J*)** NG+pECFC displayed minimal IgG extravasation and maintained strong astrocyte interaction at the cerebrovasculature. **(*I*** FGR+pECFC demonstrated less frequent IgG extravasation and significantly lower IgG-positive labelled area compared with untreated FGR **(*K;*** each dot represents individual measurements). All values are expressed as mean +/- SEM. FGR (*n*=7) vs FGR+SC (*n*=6). Two-way *ANOVA* with Tukey post-hoc test (**p<0.01, ****p<0.001). For ***K***; unpaired Student’s t-test (* *p*<0.05) (Scale bars: 50µm; for *A* & *G*: low magnification: 200µm).

Similarly, we examined permeability of the BBB with the endogenous immunoglobulin IgG which is a larger molecular weight protein. Labelling of IgG showed sporadic extravasation into the parenchyma of 71.4% of FGR brains examined with juxtavascular astrocytes in these regions displaying activated morphology but loss of end-feet contact with the vessel (Figure 3G-g’). NG and NG+pECFC groups showed limited to no IgG labelling and were therefore excluded from statistical analyses (1 animal displayed extravasation in both NG & NG+pECFC; Figure 3H & J). FGR+pECFC brains showed fewer incidences of extravasation (50%), and significantly decreased IgG areal density (Figure 3I & K).

Reduction in vessel-astrocyte interaction may be associated with less mature astrocytes at the vessel-glia interface in the FGR newborn brain. We therefore co-labelled for S100β (marker of mature astrocytes) and GFAP to investigate altered astrocyte interaction with microvessels. In NG animals, we observed an abundance of S100β-positive juxtavascular astrocytes and co-localisation with GFAP with consistent contact of end-feet with the vasculature (Figure 4A). FGR demonstrated altered labelling patterns, with significantly less S100β-GFAP positive cells along the vasculature compared with NG animals (p=0.007; Figure 4B & G) as well as an overall decrease in GFAP-positive vessel coverage (p=0.004; Figure 4E). FGR+pECFC displayed labelling similar to that observed in NG, with an abundance of S100β-GFAP-positive labelling at the vasculature (Figure 4C). FGR+pECFC showed similar GFAP coverage to that of NG brains, with a strong trend towards improving astrocyte coverage when compared with FGR (p=0.056; Figure 4E). These findings suggest that pECFC administration on first day of life promotes the maturation of juxtavascular astrocytes.

**Figure 4.**
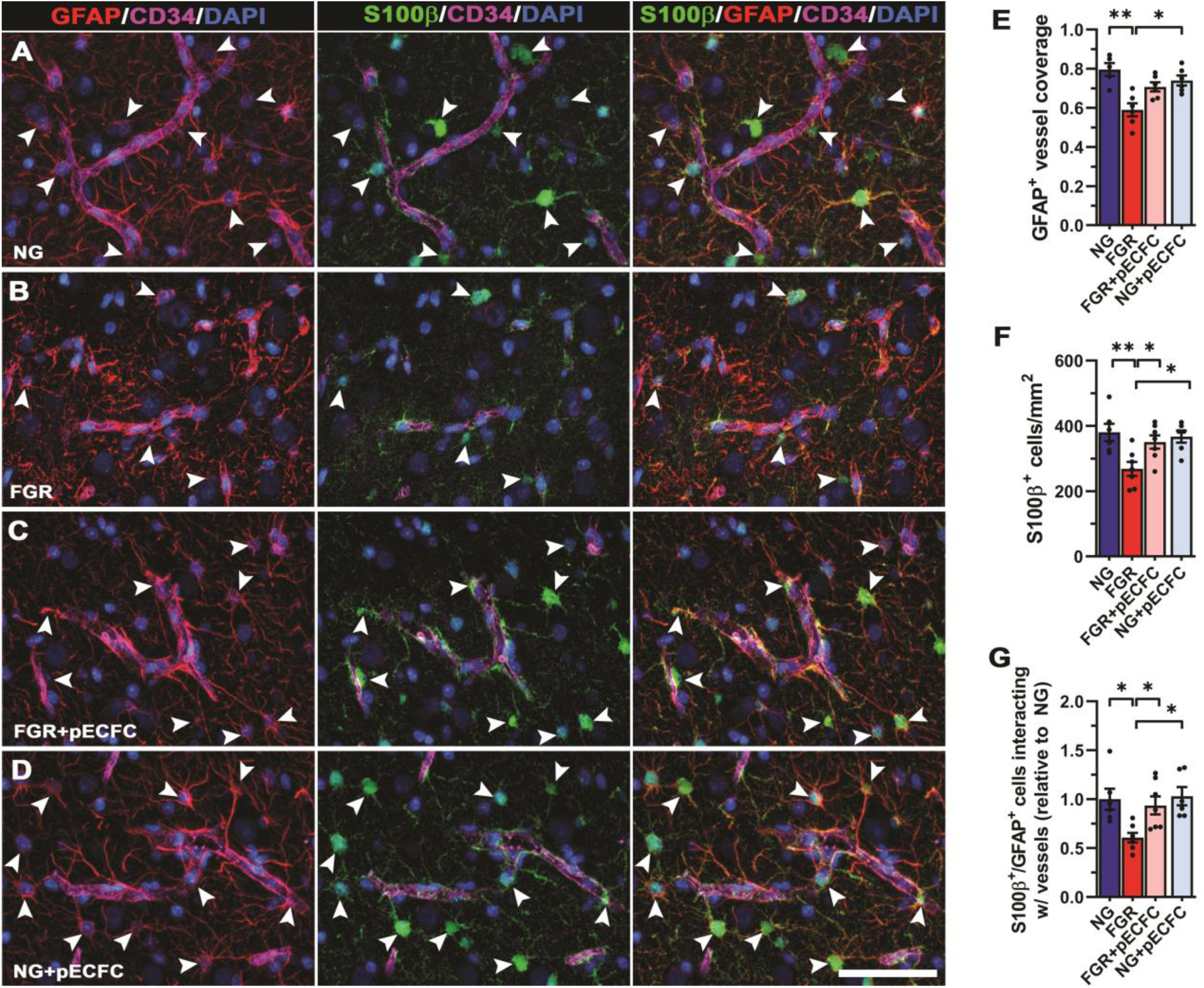
Recovery of mature astrocytes at the NVU of FGR brain following pECFC administration. Representative labelling for pan-astrocyte marker (GFAP; red) and mature astrocyte marker (S100β; green) in the cortex of pig at day 4. (**A**) NG showed strong S100β labelling co-localized to GFAP (arrowheads). These mature astrocytes demonstrate strong interaction of end-feet enveloping the length of neurovasculature.(**B**) FGR displayed intense thickened process labelling around the vasculature, indicative of reactive state. Less frequent S100β labelling was observed in the cortex as well as at the vasculature specifically. GFAP labelling along the vasculature was more sporadic and uneven when compared with NG. **(*C*)** FGR+pECFC displayed mature astrocytic labelling comparable to NG, with astrocyte end-feet displaying more contact with vasculature. **(*D*)** NG pECFC treated showed no alteration in cell count or morphology. **(E)** Co-localization analysis demonstrated a decrease in GFAP-positive vessel coverage in FGR brain compared with NG. **(F)** The reduction in S100β cell counts in FGR was ameliorated following pECFC treatment. **(G)** Quantification of S100β-GFAP positive cells interacting with the vasculature confirmed a significant decrease in FGR relative to NG. FGR+pECFC displayed similar numbers of S100β-GFAP positive cells interacting with the vasculature to that observed in NG. All values are expressed as mean +/- SEM (minimum *n*=5 for all groups). Two-way *ANOVA* with Tukey post-hoc test (*p<0.05) (Scale bars: 50µm).

### pECFC treatment attenuates glial cell activation and modifies the pro-inflammatory environment of the FGR brain

Both microglia and astrocytes are key drivers of the inflammatory response following brain injury, with the associated alterations in morphology being a hallmark to this pathway. We and others have reported early and persistent inflammation is associated with glial activation in the FGR brain (*16, 17, 35, 37, 38*). We examined if the administration of pECFC modulates the pro-inflammatory environment of the FGR brain. Labelling with the microglia marker Iba-1 revealed microglia in the parietal cortex of NG brain that displayed characteristic ramified (resting) morphology, with long fine process extensions and even distribution across the cortex (Figure 5A). In the FGR brain, microglia displayed enlarged cell bodies and thickened retracted processes as previously reported (Figure 5A) (*16, 35*) with a significant increase in the number of total microglia (p=0.0007) and activated microglia (p<0.0001) in FGR when compared with NG (Figure 5C & D, respectively).

**Figure 5.**
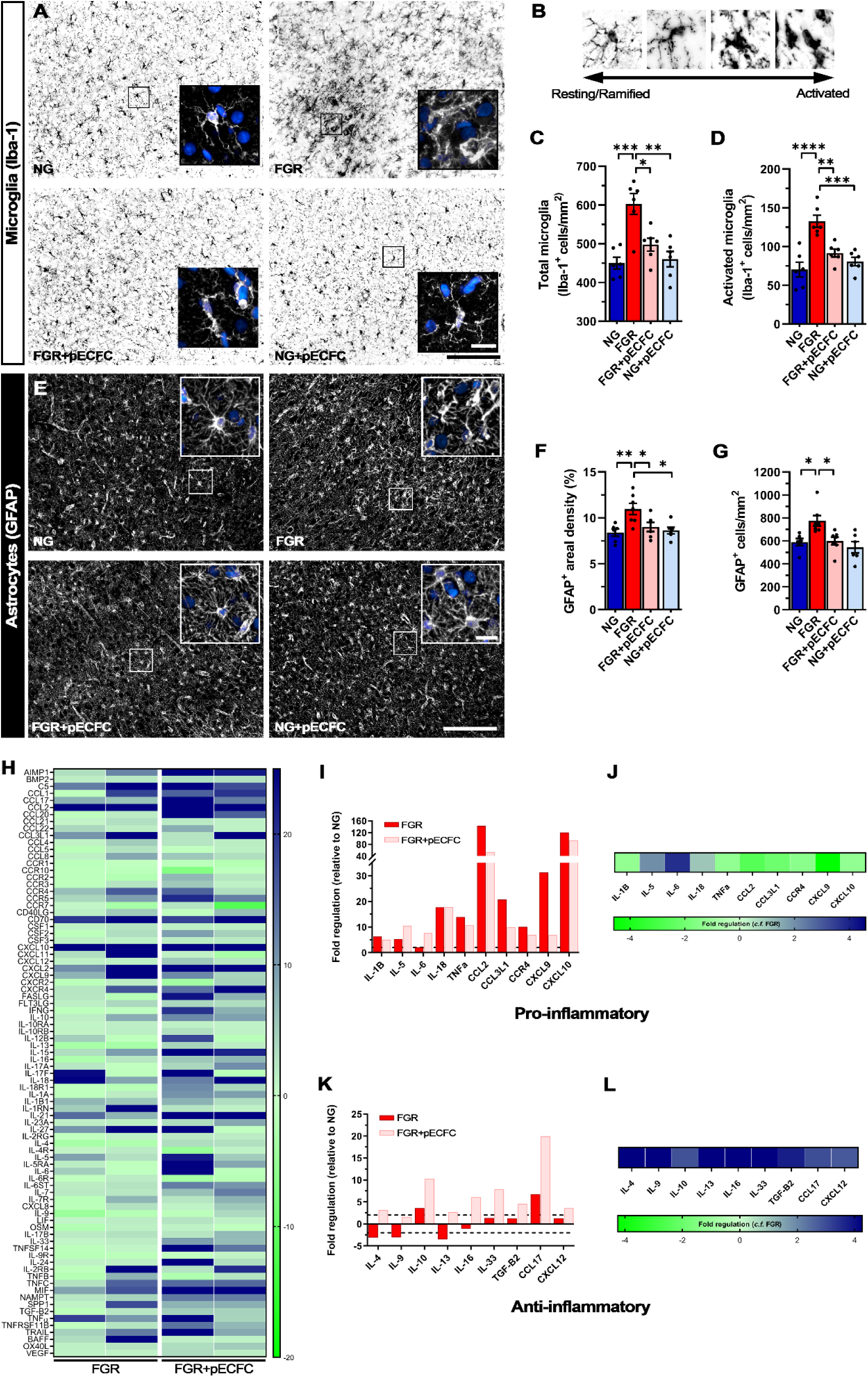
pECFC treatment attenuates glial activation in FGR brains. Representative labelling of microglial cells (Iba-1; black) in the parietal cortex. **(*A*)** NG brains display typical ramified microglial morphology, with even distribution, small cell bodies, and extended processes (see insert). FGR demonstrate significant activation of microglia with dense cellular bodies, retraction and thickening of processes, and a high degree of cellular overlap. pECFC-treated animals displayed morphology comparable to NG. **(*B*)** Microglia were assigned as resting or ramified based on well-established morphology associated with activation state of glial cells. **(*C*)** FGR demonstrated significantly higher total number of Iba-1 positive cells and activated microglia **(*D*)**. Similar trends were observed in the white matter of FGR brain (see **Supp. Figure. 3**). **(*E*)** Astrocytes (GFAP; white) showed evident alterations in morphology. NG displayed typical star like cells with long process extensions. FGR displayed activated astrocyte morphology, with retracted processes and enlargement of cell bodies. pECFC-treated animals displayed similar morphology to NG. **(*F*)** Treatment with pECFCs reduced astrocyte activation as assessed with aerial coverage of GFAP-positive stained regions. **(*G*)** pECFC treatment also reduced the number of astrocytes in the cortex toward levels comparable to other groups. **(*H*)** Heat map of porcine inflammatory cytokine and receptors arrays demonstrates altered expression of pro- and anti- inflammatory mediators in the cortex of FGR relative to NG. FGR+pECFC showed alterations in inflammatory genes relative to NG and untreated FGR. Expression of well-characterized pro- inflammatory **(*I*)** and anti-inflammatory **(*K*)** genes relative to NG. FGR+pECFC displayed down- regulation of pro-inflammatory **(*J*)** and up-regulation of key anti-inflammatory genes **(*L*)** when compared with untreated FGR. All values are expressed as mean +/- SEM (minimum n=6 for all groups). Two-way ANOVA with Tukey post-hoc test (*p<0.05, **p<0.01, ***p<0.001, ****p<0.0001) (Scale bars: 50 µm).

Treatment with pECFC in FGR resulted in microglial morphology similar to NG (resting state) with significantly lower numbers of total microglia (p<0.0001) and activated microglia (p<0.0001) compared with FGR brain (Figure 5C & D, respectively). Microglia were also examined in white matter regions including the intragyral (IGWM), subcortical (SCWM), and periventricular white matter (PVWM) (Supp. Figure 2A & B). Similar findings were observed with FGR displaying increased activation of microglia relative to NG, which was largely ameliorated in the pECFC- treated FGR animals (Supp. Figure 2A & C). No significant difference in Iba-1-positive activated microglia were evident in PVWM between pECFC-treated and untreated FGR animals (Supp. Figure 2C). Given our pECFC preparation contains MSCs and previous studies have described the anti-inflammatory properties of MSC cells we also administered MSCs alone as a comparison treatment group. MSCs alone demonstrated similar potency in the cortex, with largely normalised microglial morphology and a significant reduction in the number of activated microglia (Supp. Figure 3A & B).

We next examined astrocytes, which are involved in maintaining homeostasis of the CNS, promoting neuronal development and responding to insults. In the NG pig brain GFAP-positive astrocytes demonstrated multiple long branching processes from the cell body typical of normal astrocyte morphology and were evenly dispersed across the cortex (Figure 5E). In comparison, astrocytes in the FGR brain displayed reactive morphology, with retraction and thickening of processes into the cell body (Figure 5E) as previously demonstrated (*16*). We observed an increase in the number of GFAP-positive cells (p=0.019; Figure 5G) compared with NG indicating astrogliosis in the FGR brain. GFAP-positive astrocyte density was also significantly increased in the FGR cortex suggesting an increase in reactive astrocyte morphology (p=0.006; Figure 5F), IGWM, SCWM and PVWM (Supp Figure 2D & E) as previously demonstrated (*16*).

Both pECFC and MSC only treated FGR groups presented ramified astrocytic morphology comparable to that observed in NG animals. pECFC-treated FGR animals displayed significantly lower GFAP-positive astrocyte density in the cortex (p=0.046; Figure 5F), IGWM and PVWM (Supp Figure 2E), as well as lower cell counts (p=0.020; Figure 5G) compared with FGR. Treatment with MSCs alone did not significantly reduce GFAP positive labelling compared with FGR (Supp. Figure 3C & D). Together these findings suggest an anti-inflammatory effect modulated by MSCs but that combination with ECFC – pECFCs is more efficacious.

Using polymerase chain reaction (PCR) arrays of chemokines and cytokines, we report altered expression of both pro- and anti-inflammatory genes in the FGR cortex compared with NG brains (Figure 5H). Critical mediators of a pro-inflammatory response including CCL2, CXCL10, IL-1β and TNFα were upregulated in FGR compared with NG (Figure 5H & I). These cytokines are expressed in glial and neuronal cells in the FGR brain at postnatal day 4 (*16*) and likely contribute to the ongoing pro-inflammatory cycle. pECFC-treated FGR animals showed reductions in pro-inflammatory genes, specifically key mediators such as IL-1β, TNFα, and CXCL10 (Figure 5H- J). Anti-inflammatory mediators IL-4, IL9, and TGF-β2 showed lower expression in FGR and were upregulated following pECFC treatment in FGR animals (Figure 5H, K & L). Our findings indicate a modulation of inflammatory mediators rather than suppression or complete cessation following cell administration. Together with the reduction in overt glial activation, it is likely the inflammatory environment is more precisely regulated in pECFC-treated FGR brains when compared with FGR.

### pECFC treatment reduces neuronal apoptosis in the FGR brain

We have previously reported that FGR results in significant reductions in the number of NeuN-positive neuronal cells in the parietal cortex compared with NG at postnatal day 4 (*16*). FGR cortex demonstrated regions sparse in neurons, as labelled with the neuronal nuclei marker (NeuN) and the structural neuronal marker (MAP2) (Figure 6A). In contrast, pECFC-treated FGR brains showed densely packed neurons well distributed and organised into the cortical layers recapitulating observations in NG animals (Figure 6A). Quantification of labelling showed a significant reduction in NeuN-positive cells in FGR compared with NG (p=0.0005; Figure 6B). MAP2 labelling was also decreased in FGR compared with NG (p=0.005; Figure 6C). Following pECFC treatment in FGR, there was a recovery in the number of NeuN-positive cells (p=0.039) and MAP2-positive labelling resembled that observed in NG animals (p=0.015).

**Figure 6.**
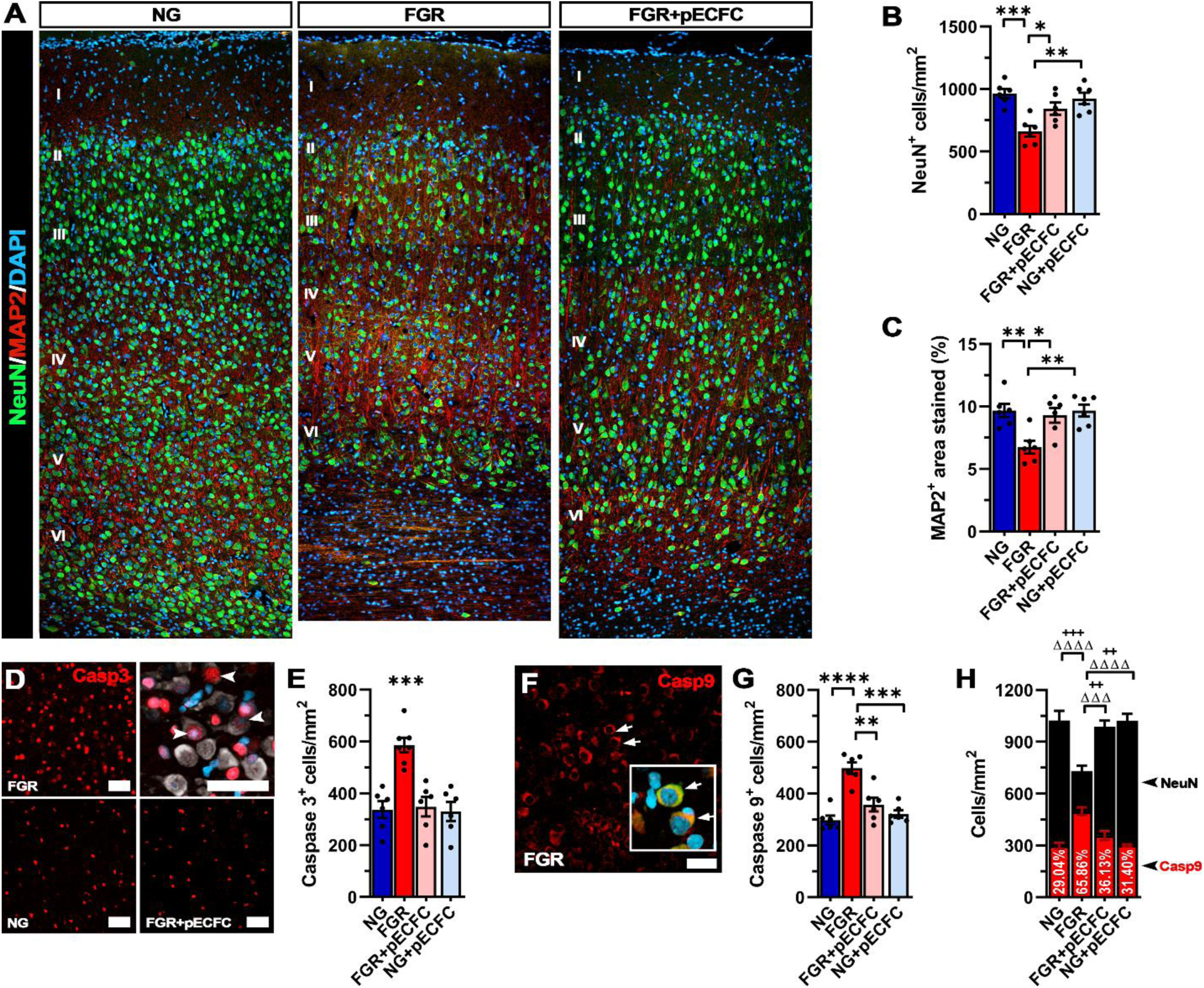
Neuronal integrity is recovered in FGR cortex following pECFC treatment. **(*A*)** Representative labelling of neuronal staining in the cortex at day 4. NG demonstrate dense labelling of mature neurons (NeuN; green) and microtubule associated protein 2 (MAP2; red). FGR demonstrate altered labelling for both NeuN (green) and MAP2 (red). Neuronal labelling is sparse and MAP2 labelling showed weaker perikarya and dendritic staining compared with other groups. These observations were evident across most cortical layers (II-VI). **(*B*)** Quantification of labelling demonstrated decreased numbers of NeuN-positive cells in FGR cortex which was not observed in FGR+pECFC brain. **(*C*)** MAP2 quantification also demonstrated a similar trend with FGR+pECFC displaying values similar to NG cortex. **(*D*)** FGR cortex demonstrated high numbers of casp3-positive cells, with limited co-localisation to mature neurons (NeuN; grey). **(*E*)** FGR displayed elevated cleaved-casp3-positive cells relative to all other groups. **(*F*)** Labelling for the initiator caspase (Caspase 9; red) showed clear localisation in mature neurons (see insert: co- localisation with NeuN indicated with arrows). **(*G*)** Quantification of casp9-positive cells showed an increase in FGR relative to all other groups. **(*H*)** Co-localisation analysis of casp9 /NeuN cells demonstrated significantly increased proportion of neurons undergoing initiation of apoptosis in FGR cortex relative to all other groups (+ denotes significance between neuronal counts, Δ significance between casp9 counts). All values are expressed as mean +/- SEM (minimum *n*=6 for all groups). Two-way *ANOVA* with Tukey post-hoc test (*p<0.05, **p<0.01, ***p<0.001, ****p<0.0001) (Scale bars: 50µm).

We have previously demonstrated an increase in cellular apoptosis in the FGR brain and the ability of the anti-inflammatory ibuprofen to ameliorate this increase (*16, 35*). We therefore proceeded to determine whether administration of pECFC could also modulate apoptotic activity in the FGR brain.

FGR brain showed a significantly higher number of cells positive for cleaved caspase-3 (Casp3) compared with NG brain (p<0.001; Figure 6D & E). Administration of pECFCs significantly reduced the number of Casp3-positive cells in the parietal cortex compared with untreated FGR animals (p<0.001; Figure 6D & E). Although some of the Casp3-positive cells were also NeuN-positive, the majority of Casp3-positive cells did not harbor this marker. We therefore investigated whether the neuronal cell bodies are undergoing early apoptosis using caspase-9 (Casp9). We observed a significant increase in Casp9-positive cells in FGR compared with NG (p<0.001; Figure 6F & G), which was ameliorated following pECFC treatment (p=0.002; Figure 6F & G). In those cells that labelled with Casp9, 65% were found to co-label with NeuN (p<0.0001) in FGR compared with only 29% in NG animals, suggesting that neuronal cells have been flagged to undergo apoptosis (Figure 6H). pECFC treatment significantly reduced the proportion of Casp9- positive apoptotic neurons in the FGR-treated animals compared with FGR animals. There was no significant differences in Casp9-positive cell counts between NG animals and NG pECFC-treated (p=0.846) animals (p=0.201).

### pECFC treatment ameliorates white matter disruption in the FGR brain

FGR is associated with significant alterations in white matter structure and organization (*39, 40*). Our previous studies have reported decreased myelination and impaired myelination based on Luxol fast blue staining of the white matter in FGR animals at postnatal day 1 and 4 (*16, 35*). Here, we investigated expression of myelin binding protein (MBP), neurofilament (NF), and the pan-oligodendrocyte marker (Olig2) in white matter. NG brains demonstrated well organised and consistent labelling for each of these markers along lengths of the white matter tracts, with strong MBP and NF co-localisation (72.3% co-localisation; Figure 7A-Ac, D). In comparison, FGR white matter presented more truncated and uneven labelling across the length of the white matter tracts for both MBP and NF (Figure 7B & B’). Areal analysis showed a significant loss in MBP-positive labelling in the white matter of FGR compared with NG (p=0.004; Figure 7E). Co-localisation analysis found evidence of NF-positive axons displaying minimal to no MBP labelling, indicating partial loss of myelination along the axonal length (FGR: 54.8% co-localisation; Figure 7Ba-c & D). pECFC-treated FGR animals displayed similar labelling patterns to those observed in NG brain (Figure 7C-Cc & D). pECFC-treated FGR animals displayed significantly higher MBP-positive labelling compared with untreated-FGR animals (p=0.009; Figure 7E). A decrease in the number of Olig2-positive cells was also observed in FGR animals compared with NG (p=0.003; Figure 7F) which was restored by pECFC treatment (Figure 7F). This decrease in Olig2 cell count was attributed to an increase in apoptosis of Olig2-postive cells. Co-labelling of Olig2 with the apoptotic marker cleaved caspase-3 (Casp3), showed a significantly higher percentage of Olig2 cells were undergoing apoptosis in FGR compared with NG (FGR: 51% cf NG: 23%, p=0.0001; Supp Figure 4A & B). Once again, pECFC-treated FGR showed a reduction in the number of apoptotic Olig2 cells (33%) compared with FGR (p=0.012; Supp Figure 4A & B). We next examined whether the loss of Olig2-postive cells in the white matter was associated with the degree of MBP labelling. Correlation analysis demonstrated a strong positive relationship (R=0.700, p<0.001; Figure 7G), of loss of Olig2 cells and lower myelination (*16*). White matter tracts in FGR brain displayed more dispersion and varied orientation based on MBP-positive labelling. Analysis of MBP orientation confirmed a higher degree of dispersion in the FGR brain compared with NG brain (p=0.015; Supp Figure 4C & D). pECFC-treated FGR animals displayed significantly lower WM tract dispersion than FGR animals (p=0.046; Supp Figure 4C & D). These findings indicate administration of pECFCs promoted Olig2 survival, maintaining axonal myelination and organisation of white matter tracts.

**Figure 7.**
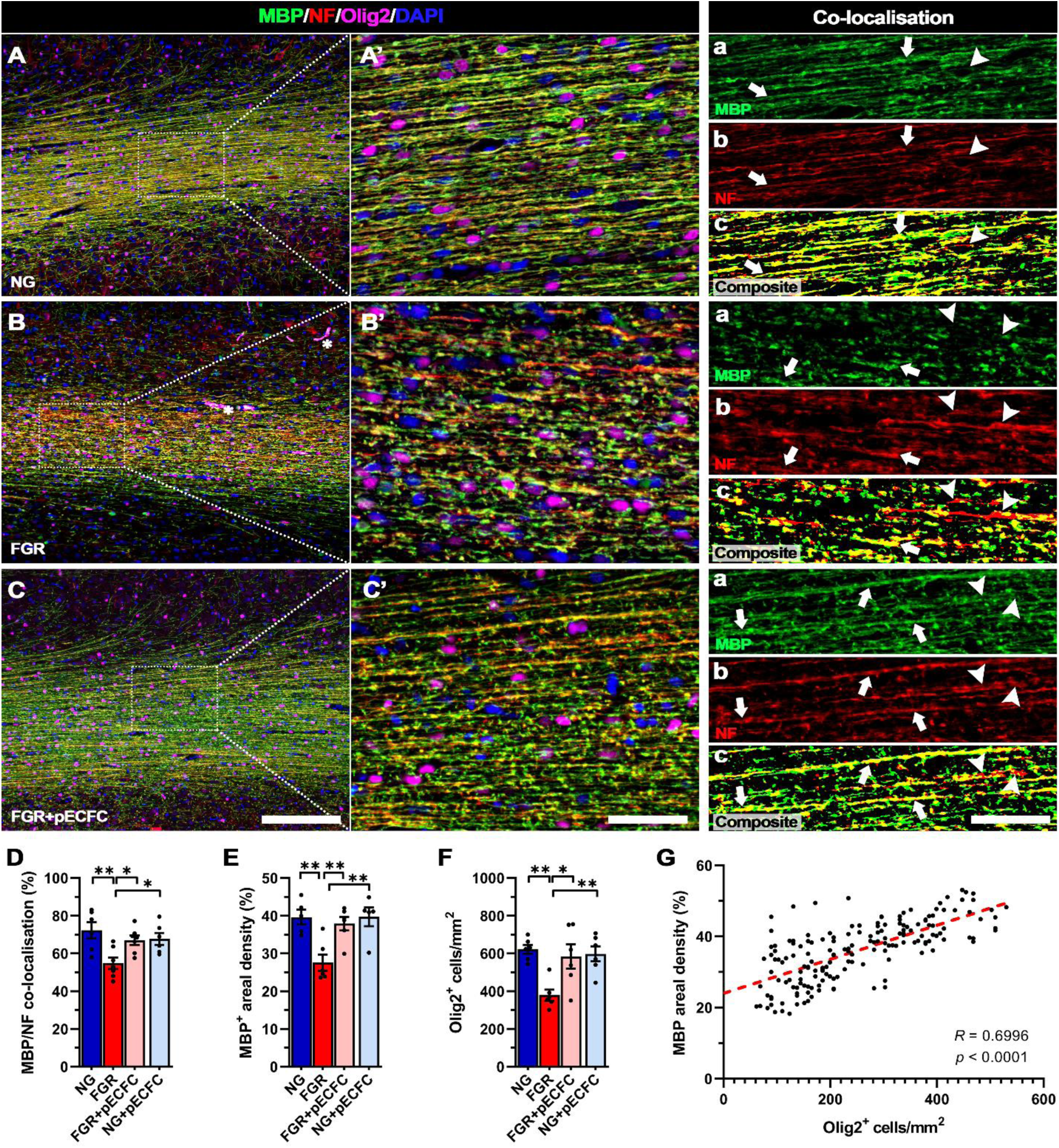
Improved axonal myelination following pECFC treatment in FGR. Representative labelling of myelin (MBP), neurofilament (NF) and pan-oligodendrocyte (Olig2) expression in white matter of NG, FGR and FGR+pECFC brains at postnatal day 4. (***A***) NG showed robust labelling for each marker with consistent presence of Olig2-positive cells along fibres. MBP and NF displayed strong co-localisation and structure throughout the white matter (***A’a-A’c***; arrows). (***B***) FGR display disrupted labelling of MBP and NF, with decreased Olig2-positive cells. There was observable disruption in labelling patterns with an evident loss in co-localisation of MBP to NF (***B’***; arrowheads). (***C***) FGR+pECFC brains showed normalised labelling and organisation of MBP and NF similar to that observed in NG. (***D***) Analysis of MBP/NF co-localisation in white matter shows a loss in axonal myelination in FGR compared with all groups.(***E***) Quantification of MBP areal density (%) showed significantly decreased areal coverage and (***F***) decreased Olig2- positive cells/mm^2^ in FGR white matter relative to all groups, both of which were largely ameliorated following pECFC treatment. (***G***) Correlative analysis demonstrates a positive relationship between Olig2-positive cell count and MBP areal coverage. All values are expressed as mean +/- SEM (minimum *n*=6 for all groups). Two-way *ANOVA* with Tukey post-hoc test (*p<0.05, **p <0.01, ***p<0.001) (Scale bars: A-C: 200µm; A’-C’ & a-c: 50µm).

## Discussion

Neurodevelopmental delays occur in 24-53% of FGR infants at 2 years of age (*41, 42*). There are no treatment options currently available to protect the FGR newborn. This is the first study to provide evidence that postnatal intervention of placentally derived pECFC treatment affords neuroprotection in the FGR piglet. We demonstrate pECFC infusion improves vascularization, NVU integrity, recovery of neuronal maturation and white matter development in the FGR brain. Our findings indicate that pECFC treatment provides neuroprotection in the FGR neonate via targeting both inflammation and the NVU.

### pECFC treatment promotes vessel density and neurovascular unit integrity

In the current and previous study, we observed changes to neurons and glial cells in the postnatal FGR brain (*16, 35*). Examination of vascular endothelial cells and basement membrane revealed a significant reduction in these cell types suggesting a loss in cerebrovasculature in the FGR brain with a significant decrease in total vascular length and vessel branching in the FGR brain.

MSC only treatment had no significant effect on improving the vasculature in the FGR brain. However, pECFC treatment not only increased vessel density it also improved vascular length in the FGR brain. pECFC treatment did not show a significant difference in vessel branching suggesting pECFC treatment does not promote unwarranted angiogenesis. These findings support the potential of pECFCs to improve cerebrovascularisation in the FGR neonate. On the contrary to our findings of reduced endothelial cells in FGR brain, an increase in endothelial cells (GLUT-1) were reported in both FGR and FGR umbilical cord blood (UCB) treated lambs in comparison to normally grown lambs (*23*). This discrepancy may be due to the marker used to label the endothelial cells. Yet, Grandvuillemin *et al*., 2017 showed improved capillary density (eNOS) follow both UCB and ECFC treatment at 7 days and 12 weeks after treatment in a neonatal hypoxic-ischemic encephalopathy (HIE) rat model (*43*). The authors also showed an increase in cerebral blood flow at 12 weeks for both treatments demonstrating evident functional improvement to the cerebral vasculature.

There was evidence of BBB disruption in the piglet FGR brain, with endogenous proteins albumin located in the perivascular space and extravasation of IgG into the brain parenchyma. Juxtavascular astrocytes displayed activated morphology including hypertropic end-feet, and in some instances we found absence of vessel-glial contact. pECFC treatment improved BBB integrity with low intensity albumin labelling restricted to the vessel lumen and juxtavascular astrocytes reverting to a ramified astrocyte morphology. Even though pECFC treatment did not significantly reduce the percentage of vessels with perivascular albumin labelling it did significantly reduce albumin areal density compared with FGR. In a lamb FGR model, albumin was shown to enter the tissue parenchyma which we did not observe in our current study, however the authors also showed a reduction in albumin extravasation following UCB treatment supporting the effect of stem cells on BBB integrity (*23*).

Impaired astrocyte end-feet interaction with neurovasculature may be associated with altered NVU integrity in the FGR brain (*23*). Following pECFC treatment, the FGR piglets displayed similar characteristics to NG animals with an abundance of juxtavascular astrocytes in consistent contact along the vasculature. This combination of ECFCs and MSCs may be working together to stabilize the NVU and by reducing the pro-inflammatory environment which in turn results in juxtavascular astrocytes returning to their normal function at the NVU. In the FGR lamb, while UCB treatment did not alter astrogliosis, the authors observed an increased association of smooth muscle proteins of the basal lamina with pericytes in the NVU (*23*).

### pECFC treatment attenuates glial activation in the FGR piglet brain

Clinical and animal studies provide evidence that inflammation is perpetuated long after birth, providing the opportunity of targeting inflammation soon after birth which may be critical to reducing detrimental inflammatory events and subsequent injury to the FGR brain (*44, 45*). We and others have reported that early and persistent inflammation in the brain is associated with glial activation in the FGR neonate (*13, 16, 23, 35*). In the current study we have demonstrated an overt glial response in the FGR piglet brain after birth with an increase in activated microglia and astrocytes in both the grey and white matter. Microglia responded to both MSC only treatment and pECFC treatment with a decrease in both the number and activation of microglia throughout the brain parenchyma. However, pECFC treatment was more efficacious than treatment with MSC alone at minimizing astrocyte activation in the FGR piglet brain. The MSC alone treatment results are corroborated by an FGR lamb study (*23*). The authors demonstrated when allogeneic umbilical cord blood (UCB) mononuclear cells (25 million/kg) was administered 1h after birth, activated microglial cells were reduced but not astrocytes in the white matter 24 hours post-treatment (*23*). An FGR rat study also showed that umbilical cord derived mesenchymal stromal cells (UC-MSC) (1x10^5^) did not affect the number of astrocytes 7 days after treatment (*46*). Astrocytes have many functions in the brain and are located in the brain parenchyma as well as at the NVU. The positive response of glia in the tissue parenchyma we observe following pECFC treatment may be due to restoration of cerebrovascularisation and NVU integrity. As the combination of ECFCs and MSCs are stabilizing the NVU and increasing the anti-inflammatory state (discussed below), these interrupt the perpetual inflammatory environment in the brain. However, one study showed a similar inflammatory response between UBC and ECFC treatment in a neonatal HIE rat model (*43*). This study administered stem cells intraperitoneally 48h after insult - 10^5^ ECFC or 10^7^ UCB. At 7 days post treatment, they demonstrated a reduction in neuroinflammation (iNOS) after both treatments, but no change in astrocyte numbers. However, at 12 weeks a significant reduction in GFAP-positive cells was reported in both groups demonstrating a delayed decrease in astrogliosis. Demonstrating long-term studies are necessary to determine the longevity of treatment effects.

Increased glial cell activation is associated with a pro-inflammatory environment in the FGR brain (*44*). The current study concurs with previous reports showing an increase in pro-inflammatory cytokines in the FGR brain and an altered anti-inflammatory profile (*16, 35*). We have previously demonstrated anti-inflammatory treatment (ibuprofen) in the FGR newborn piglet reduces levels of pro-inflammatory cytokines (*16*), however in the current study pECFC treatment exerted a greater effect on increasing anti-inflammatory cytokines. The prominent increases in anti-inflammatory cytokines may be due to reprograming of the microglia into an anti-inflammatory state. Recent *in vitro* studies suggest that a key therapeutic function of MSCs is their ability to reprogram brain microglia into an anti-inflammatory state characterized by increased phagocytic activity and upregulated expression of anti-inflammatory mediators, which in turn contributes to reduced inflammation and promotes tissue repair (*47*). These findings are corroborated in an FGR rat study examining UC-MSC treatment (*46*). The authors demonstrate no significant effect of treatment on pro-inflammatory microglia however anti-inflammatory microglia were significantly increased in the treatment group compared with sham, demonstrating an anti-inflammatory effect of stem cell treatment in the FGR rat. An FGR lamb study reported a reduction in the pro-inflammatory cytokine TNFα following UCB treatment, similar to the response we demonstrated following pECFC treatment. However, they report no effect of UCB on any other pro-inflammatory cytokines measured (*23*). We demonstrated that pECFCs have a combined effect of reducing pro-inflammatory cytokines while concurrently increasing anti-inflammatory cytokines. MSCs and ECFCs target different cell populations to maintain a healthy brain environment, and therefore this multicellular effect may be the key to modulating the inflammatory state in the brain.

### Neuronal and white matter integrity is recovered following pECFC treatment in the FGR piglet brain

Clinical imaging studies describe persistent grey and white matter disruption in the human FGR neonate that are associated with developmental disabilities at 1 year of age (*39, 48*). Ongoing neuronal and white matter impairment are also evident in the FGR piglet brain after birth at a microscopic level (*16, 34*). In the current study we saw regions sparse in neurons (mature and structural neurons) and disruption to mature myelin and oligodendrocytes throughout the white matter in the FGR brain. Inflammation is a key driver of neuronal and white matter injury and when postnatally targeted with an anti-inflammatory intervention as we have done here with pECFCs and previously with the anti-inflammatory ibuprofen, a corresponding reduction in neuronal and white matter disruption is evident (*16*).

The loss in neurons observed in the FGR brain may be attributed to pro-apoptotic events as a result of the pro-inflammatory environment. We reported a significant increase in the early apoptotic initiator, caspase-9 in neurons in the FGR brain. We have previously demonstrated an increase in cellular apoptosis and the ability of the anti-inflammatory ibuprofen to ameliorate this increase (*35*). pECFC treatment reduced neuronal cells that labelled for both caspase-3 and caspase-9. The modulation of the inflammatory response following stem cell administration likely contributed, either directly or indirectly, to the reduced apoptotic activity in the FGR brain, An HI rat study reported similar reduction in apoptotic cells (TUNEL) and concurrent recovery to neuronal cell counts 7 days after treatment with either UCB or ECFCs (*43*). This recovery to neurons could be due to reversal of apoptosis or reinstatement of normal development.

We show not only disruption to myelination as previously demonstrated (*16, 34*), but also disruption to oligodendrocytes as observed in other animal models of FGR (*14, 17, 49, 50*). We demonstrate partial loss of myelination along the axon in the FGR brain with increased apoptotic oligodendrocytes. White matter tracts in the FGR brain displayed more dispersion and varied orientation suggesting that loss of oligodendrocytes in FGR results in decreased and disorganised myelination of white matter tracts which likely contributes to the long-term white matter alterations reported in the FGR brain (*51*). Administration of pECFC treatment promoted oligodendrocyte survival, maintaining axonal myelination and organisation of the white matter tracts. Only one other study has examined white matter response to stem cell treatment in the FGR newborn. This study in the FGR rat showed no difference in white matter volume following UC- MSC treatment (*46*). However, this novel FGR model may not be appropriate to study FGR with no detectable white matter injury. White matter injury is a hallmark feature of FGR (*52*) as such there should be noticeable white matter changes in any FGR animal model.

## Conclusion

We regard pECFC treatment as a promising multicellular approach to protecting the FGR newborn brain. MSC therapy alone has variable neuroprotective effects through modulation of inflammation in small animal models of cerebral ischemia (*53–55*). However with the addition of ECFCs to aid in vascular regeneration, this primed stem cell preparation has the potential to significantly improve brain outcomes for FGR neonates. Previous studies have shown pECFCs are not rejected in immunocompetent animal studies (*27*) demonstrating important findings where these cells can be used in allogeneic situations. The use of healthy placentas and in an off-the-shelf allogenic scenario would ensure continued availability and provide a more appropriate solution than using autologous cells from a pathologic FGR placenta (*56*). Future preclinical studies to confirm long-term efficacy and safety are essential in the success of human clinical trials for stem cell treatment in the FGR newborn.

## Materials and Methods

### Animals

Approval for this study was granted by The University of Queensland Animal Ethics Committee (UQCCR/420/17) and was carried out with respect to the National Health and Medical Research Council guidelines (Australia) and ARRIVE guidelines. Animals care was in accordance with institutional guidelines.

Newborn Large White piglets (n=36; <18 h) were collected from the UQ Gatton Piggery monitored and cared for at the Herston Medical Research Centre (HMRC) until day of euthanasia on postnatal day 4 (P4). Litter matched pairs were obtained from multiple sows (n=14 litters). Piglets were divided into five experimental groups with pigs randomly assigned to treatment groups: NG (n=8), FGR (n=8), FGR+pECFC (n=8); FGR+MSC (n=5), and NG+pECFC (n=7); with equal males and females in each group. FGR piglets were defined by birth bodyweight (<10th percentile on the day of birth) and confirmed through brain to liver weight ratio (B:L) ≥ 1 assessed post-mortem at P4 (*34, 57, 58*).

On P4, piglets were euthanised via an intraperitoneal injection of sodium phenobarbital (650mg/kg; Lethabarb, Virbac, Australia). Brains were transcardially flushed with phosphate-buffered saline, and brain tissue was collected, weighed, hemisected and coronally sliced. The right hemisphere sections were immersion fixed in 4% paraformaldehyde as previously described (*59*). The parietal cortex from the left hemisphere was snap frozen in liquid nitrogen and stored at -80 °C for mRNA analysis.

### Fluorescence-activated Cell Sorting

Primary conjugated antibodies were used for fluorescence-activated cell sorting (FACS). Placental tissues were processed, and single-cell suspension was prepared as reported previously (*30, 33*). The isolated placental CD34+ single cell suspension was incubated with human CD34- phycoerythrin (PE) (Bio-Rad; dilution 1:25), human CD45-FITC (BioLegend; dilution 1:25) and human CD31-V450 BD Biosciences: dilution 1:30) for 20min at 4 °C. Cells were flow sorted using FACSAria Fusion (Becton Dickinson). Cell doublets were removed and 7-amino-actinomycin D (7AAD) (BD Pharmingen; dilution 1:40) was used to exclude dead cells. Fluorescence minus one (FMO) control was used in gating the population of interest. To remove any remaining contaminating CD45^+^ cells from the hematopoietic lineage, only CD45^-^CD34^+Hi^ population was gated and further sorted according to their level of CD31 expression. The population of interest, the CD31^-^ and CD31^Int^ populations were sorted directly into 100% fetal bovine serum (FBS; Gibco) before being plated for cell culture.

### Stem Cell Culture

Human placental fetal ECFC and MSC were isolated using our previously published protocol (*30*). The use of human tissue was granted by the human ethics boards of The University of Queensland and the Royal Brisbane and Women’s Hospital. The isolated fetal ECFC and MSC were cultured on rat tail collagen coated tissue-culture flasks in Endothelial Growth Medium (EGM2) (Lonza) with 10% of FBS. For *in vivo* stem cells treatment, 10^6^ ECFC 10^6^ MSC were injected intravenously into NG and FGR piglets (Figure 1A & B). In a further subset of FGR animals 10^6^ MSC only were injected.

### Magnetic resonance imaging

We used magnetic resonance imaging (MRI) techniques to determine whether we could detect *in vivo* brain alterations in the FGR piglet at postnatal day 4. Piglets were anesthetised with isoflurane (1-3 %) mixed with oxygen (2L/min) for the duration of the scanning protocol. Each piglet was placed in a 300mm bore 7T ClinScan MR scanner (Bruker, Germany), running Siemens VB17. A 150 mm ID MRI rf coil was used to acquire the dynamic images of sagittal, coronal and axial slices. Images were obtained at TR/TE 8500/60ms, 1.6mm slices thickness, 64x64 acquisition matrix. Region of interest over the parietal cortex was defined on the T2 map and applied to the ADC map and raw values extracted.

### Magnetic resonance spectroscopy acquisition and analyses

1H-MR spectroscopy were obtained using the 7T Bruker/Siemens whole-body. A single spectrum was acquired the frontoparietal region of the brain from a 10 mm3 voxel using single-voxel spectroscopy with the following parameters: TR=6000 ms, TE = 60 ms and 128 averages. Metabolite spectra were exported, processed, and analysed using the Advanced Method for Accurate, Robust and Efficient Spectral Fitting (AMARES) tool within jMURI and manually phased (*60*). Before quantification, apodisation (10 Hz) was applied to the N-acetyl aspartate (NAA), creatine (Cr) and choline (Cho) peaks with Lorentzian peaks. Peak areas of metabolites were estimated using Lorentzian profiles and a priori knowledge with soft constraints (initial chemical shift, amplitude, and peak widths). NAA was set as a reference at 2ppm chemical shift. Peak area ratios were calculated for NAA/Lac, NAA/Cho, NAA/Cr, Lac/Cr, Lac/Cho, Lac/NAA and Cho/Cr.

There were no overt differences in brain structure as assessed with T2 and ADC analysis between FGR and NG animals. There were no significant MRS alterations in metabolite levels between NG and FGR. Administration of pECFCs did not result in any noticeable changes when compared with NG and FGR (Supp. Fig. 5).

### Immunohistochemistry

Brain slices containing parietal cortex from the right hemisphere (NG=8; FGR=8; FGR+pECFC=6; NG+pECFC=6; FGR+MSC=5) were embedded in paraffin and sectioned at 6µm (Pig stereotaxic map, A 5.5mm; (*61*)). For vessel structure and glial interaction analyses brain sections of 12µm thickness were used. Sections were affixed to Menzel Superfrost Plus adhesive slides and dried overnight at 37ᴼC. All sections were dewaxed and rehydrated using standard protocols followed by heat induced epitope retrieval with 10mM citrate buffer (pH 6) or TRIS-EDTA buffer (pH 9) at 90ᴼC for 20 min before cooling to room temperature (RT). Sections were blocked with 5% donkey serum in PBS with 0.5% Triton-X 100 for 1h at RT. Primary antibodies (supplementary Table 1) were incubated overnight at 4ᴼC. Slides were washed in tris-buffered saline followed by incubation with species-specific secondary fluorophores (Supp. Table 1) at RT for 1h. Sections were washed, counterstained with 4′,6-diamidino-2-phenylindole (DAPI), and mounted with Prolong Gold antifade (Molecular Probes, Invitrogen Australia, Victoria, Australia). Negative control sections without primary antibodies were processed in parallel and immunolabelling was conducted in triplicates for all animals.

### Image acquisition and analysis

Analysis of immunolabelled sections was performed using a Zeiss Axio Microscope with an Axiocam503 camera. Four pictomicrographs of the parietal cortex and white matter regions were captured for analysis from each section. For all markers, replicates were conducted with sections separated by at least 70µm. All imaging and analyses were conducted under blind conditions (KKC, JAW, ET, and LJ). Microglia were manually counted and categorised with respect to morphology as per previous studies (*16, 35*). Density analysis, co-localisation and vessel coverage analyses was undertaken using the threshold function with moments plugin in FIJI (ImageJ; Image Processing and Analysis in Java; National Institutes of Health, Bethesda, MD, USA).

### Quantitative Polymerase Chain Reaction (qPCR)

qPCR was conducted as previously reported (*16, 35*). Total RNA from frozen parietal cortex samples was isolated using RNeasy Tissue Mini Kit (Qiagen). Yield and quality were determined using a NanoDrop spectrophotometer (ND-1000 system). Reverse transcription kits (RT2 First Strand Kit; Qiagen) were used for cDNA synthesis. cDNA was pooled for each experimental group giving equal concentrations from every animal in the pooled sample. Pooled cDNA was combined with RT2 SYBR Green qPCR Mastermix (Qiagen) and loaded into the Pig Inflammatory Cytokines & Receptors RT2 Profiler^TM^ PCR Array (Qiagen, Hilden, Germany). The qPCR reactions were performed using a Qiagen Rotor-Gene Q real-time cycler. The amplified transcripts were quantified with the comparative CT method using actin, gamma 1 (ACTG1) mRNA expression levels for normalization. The same CT threshold value was used across all arrays to allow comparison between runs. Each experimental group was run across two arrays with samples randomly assigned to each array.

### Statistics

Unless otherwise specified, Two-way ANOVA with the Tukey post-hoc analyses were used to determine differences between NG and FGR animals under non-treated and stem cell treated conditions (Graph Pad Prism 9.0 software, San Diego, California, USA). Results are expressed as mean ± SEM with statistical significance accepted at p<0.05. Indicated sample sizes (n) represent biological replicates including individual samples, and have been listed in the figure legends where appropriate.

## Acknowledgements

We would like to thank the women at the Royal Brisbane and Women’s Hospital for kindly donating their placentas for this study. We would also like to thank the Flow Cytometry facility at the Translational Research Institute for their kind assistance and John Luff for his assistance with the animal experiments. **Funding:** JP salary was supported by the ARC DECRA Fellowship (DE180100984) funded by the Australian Government. KK salary was supported by the NHMRC Career Development Fellowship (APP1125290) funded by the Australian Government. A Royal Brisbane and Women’s Hospital Research Grant, Financial Markets Foundation for Children (2018-043), and UQ Early Career Researcher grant supported this work. Funding bodies did not influence the design of the study nor collection, analysis, interpretation of data, and drafting of the manuscript. **Author contributions:** KC was responsible for all brain laboratory aspects of the project, data analysis, interpretation, writing and editing the manuscript. JP was responsible for stem cell preparation, obtaining funding, critical revision, and editing the manuscript. TB was involved in obtaining funding, critical revision, and editing the manuscript. SLS was responsible for stem cell isolation and preparation, and editing the manuscript. SM was involved in animal experimentation, and editing the manuscript. ET was involved in animal experimentation, MRI analysis, and editing the manuscript. LJ was involved in animal experimentation, MRI analysis, and editing the manuscript. JS was involved in stem cell isolation and preparation, and editing the manuscript. PC was involved in obtaining funding, critical revision, and editing the manuscript. KK was involved in obtaining funding, critical revision, and editing the manuscript. JW was involved in obtaining funding, experimental design, conducting animal experiments, acquiring data, critical revision, and writing the manuscript. **Competing interests**: Authors have no competing interests. **Data and materials availability:** All data associated with this study are available in the main text or the supplementary materials.

## Definitions

FGR: fetal growth restriction
NG: normally grown
NVU: neurovascular unit
MSC: mesenchymal stem cells
ECFC: endothelial colony forming cells
pECFC: primed endothelial colony forming cells
BBB: blood brain barrier
UCB: umbilical cord blood
IGWM: intragyral white matter
SCWM: subcortical white matter
PVWM: periventricular white matter
GFAP: glial fibrillary protein
Iba-1: ionized binding protein-1
NeuN: neuronal nuclei
MAP2: microtubule-associated protein 2
IL-1β: interleukin-1β
TNFα: tumor necrosis factorα
PCR: polymerase chain reaction
HIE: hypoxic-ischemic encephalopathy
MRI: magnetic resonance imaging
NF: neurofilament
Olig2: oligodendrocyte maker 2

---

**Supplementary Table 1:**

| **Primary Antibody** | **Host** | **Dilution** | **Catalogue#** |
| --- | --- | --- | --- |
| Anti-human CD34-PE |  | 1:25 | Bio-Rad (MCA1578PE) |
| Anti-human CD45-FITC |  | 1:25 | BioLegend (304006) |
| Anti-human CD31-V450 |  | 1:30 | BD Biosciences (561653) |
| 7AAD |  | 1:40 | BD Pharmingen (559925) |
| Lamin A/C | Rabbit | 1:200 | Abcam (ab108595) |
| Vimentin | Mouse | 1:500 | Merck (CBL202) |
| Endothelial cells (CD34) | Goat | 1:1000 | R&D Systems (AF3890) |
| Ionised calcium binding adaptor molecule-1 (Iba-1) | Goat | 1:1000 | Abcam (ab5076) |
| Glial fibrillary acidic protein (GFAP) | Mouse | 1:1000 | Sigma (G3893) |
| Glial fibrillary acidic protein (GFAP) | Rabbit | 1:2000 | DAKO (Z0334) |
| Neuronal Nuclei (NeuN) | Rabbit | 1:1000 | Abcam (ab177487) |
| Microtubule-associated protein 2 (MAP2) | Mouse | 1:500 | Sigma-Aldrich (M4403) |
| Collagen IV (Col IV) | Rabbit | 1:1000 | Abcam (ab6586) |
| Cleaved Caspase-3 | Rabbit | 1:500 | Cell Signaling (#9661) |
| Caspase-9 | Rabbit | 1:500 | Abcam (ab32539) |
| Myelin Binding Protein (MBP) | Rat | 1:1000 | Abcam (7349) |
| Neurofilament (NF) | Mouse | 1:500 | Abcam (ab134306) |
| Oligodendrocyte marker 2 (Olig2) | Rabbit | 1:1000 | Genetex (GTX132732) |
| Albumin | Sheep | 1:1000 | Abcam (ab8940) |
| IgG | Goat | 1:1000 | JIR (114-005-003) |
| S100B | Rabbit | 1:1000 | DAKO (Z0311) |
| **Secondary Antibody** |  |  |  |
| α-Rat Alexafluor 488 | Donkey | 1:1000 | Invitrogen (A-21208) |
| α-Rabbit Alexafluor 488 | Donkey | 1:1000 | Invitrogen (A-21206) |
| α-Goat Alexafluor 568 | Donkey | 1:1000 | Invitrogen (A-11057) |
| α-Mouse Alexafluor 568 | Donkey | 1:1000 | Invitrogen (A10037) |
| α-Rabbit Alexafluor 594 | Donkey | 1:1000 | Invitrogen (A-21207) |
| α-Goat Alexafluor 647 | Donkey | 1:1000 | Invitrogen (A-21447) |
| α-Mouse Alexafluor 647 | Donkey | 1:1000 | Invitrogen (A-31571) |
| DAPI (4',6-Diamidino-2-Phenylindole, Dihydrochloride) | - | 1:2000 | Invitrogen (D1306) |

**Supplementary Fig. 1.**
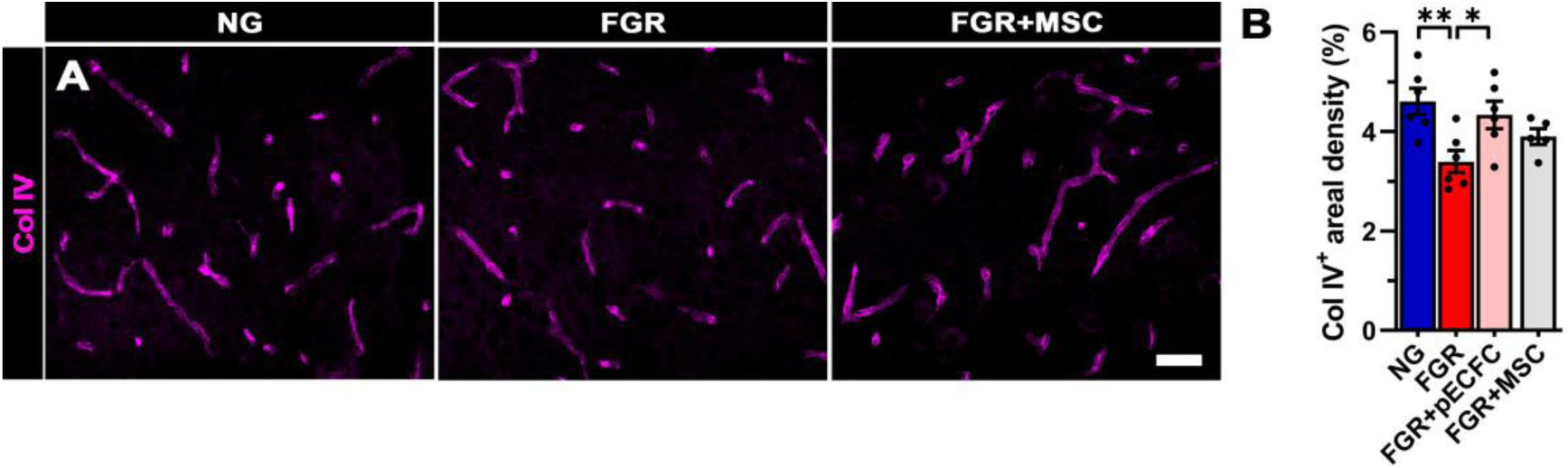
Mesenchymal stem cell treatment does not improve vascular expression of Col IV in FGR brain. (***A***) Staining for Col IV showed no significant improvement in microvessel density following treatment with MSC only. (***B***) Areal analysis confirmed no significant recovery in Col IV staining. All values are expressed as mean +/- SEM (minimum n=5 for all groups). Two-way *ANOVA* with Tukey post-hoc test (*p<0.05, **p<0.01) (Scale bars: 50 µm)

**Supplementary Fig. 2.**
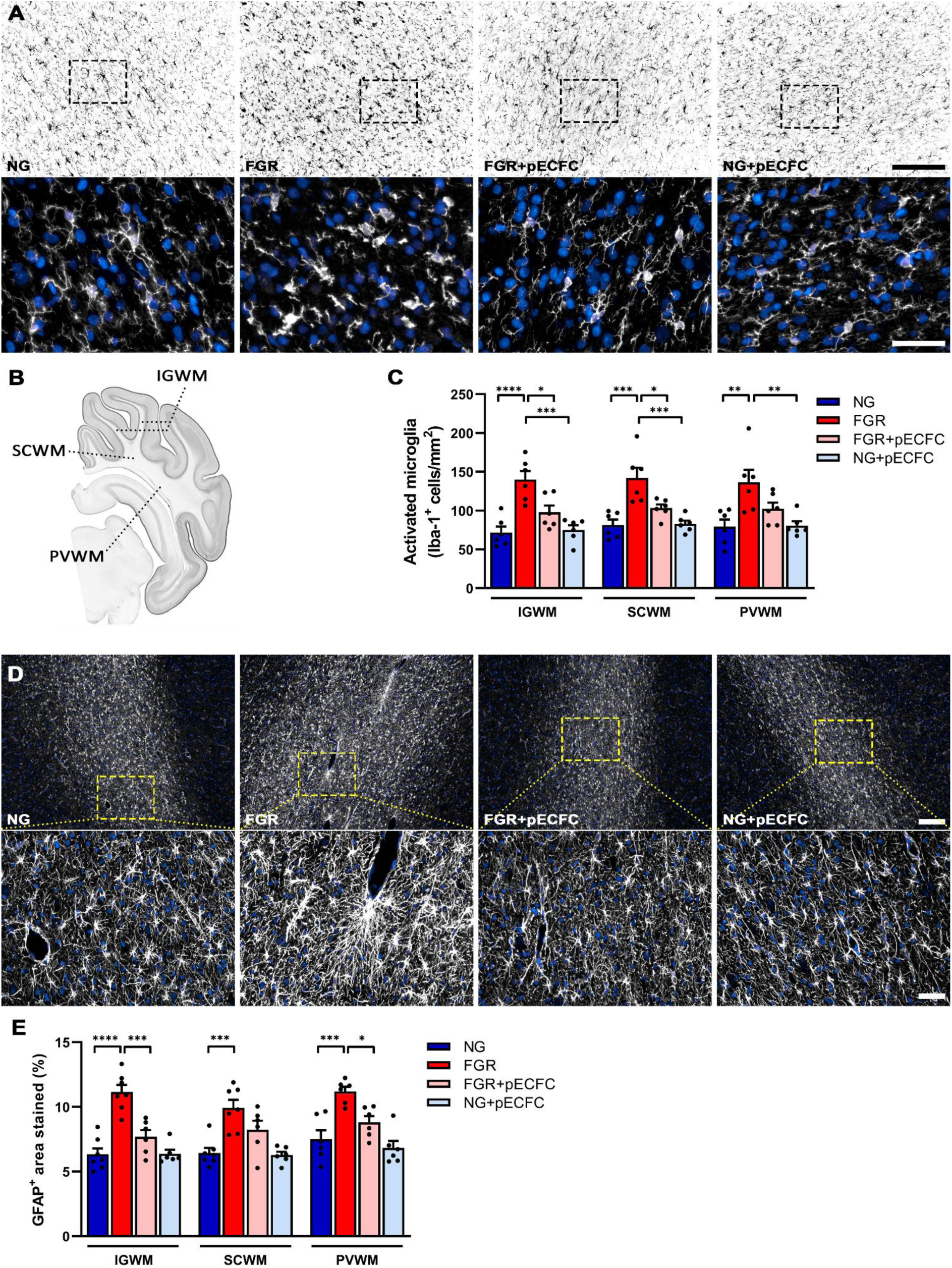
Elevated glial activation in white matter is ameliorated following pECFC treatment in FGR. (***A***) Representative staining of microglia (Iba-1) in the intragyral white matter (IGWM) of NG brain shows even cellular distribution orientated longitudinally with extended processes, indicative of ramified (resting) microglia. Microglia in FGR white matter demonstrate activated morphology. FGR+pECFC displayed normalised microglia morphology comparable to NG brains. NG+pECFC animals showed no evident alterations in microglia morphology. (***B***) Schematic outline of key white matter regions examined: IGWM, subcortical white matter (SCWM), and periventricular white matter (PVWM). (***C***) FGR brain demonstrated significantly elevated activated microglia counts in all white matter regions when compared with NG brains. Administration of pECFCs significantly ameliorated microglial activation in the IGWM and SCWM but not the PVWM. (***D***) Representative staining of astrocytes (GFAP) in the intragyral white matter (IGWM) of NG brain shows even cellular distribution of ramified (resting) astrocytes. Astrocytes in FGR white matter demonstrate activated morphology. FGR+pECFC displayed normalised astrocyte morphology comparable to NG brains. Administration of SC to NG animals showed no evident alterations in astrocyte morphology. (***E***) Areal coverage of GFAP staining was significantly elevated in FGR for all white matter regions when compared with NG brains. Administration of pECFCs significantly ameliorated astrocyte activation in the IGWM and PVWM. All values are expressed as mean +/- SEM (minimum *n*=6 for all groups). Two-way *ANOVA* with Tukey post-hoc test (*p<0.05, ***p<0.001, ****p<0.0001) (Scale bars: low magnification: 200µm; high magnification: 50µm).

**Supplementary Fig. 3.**
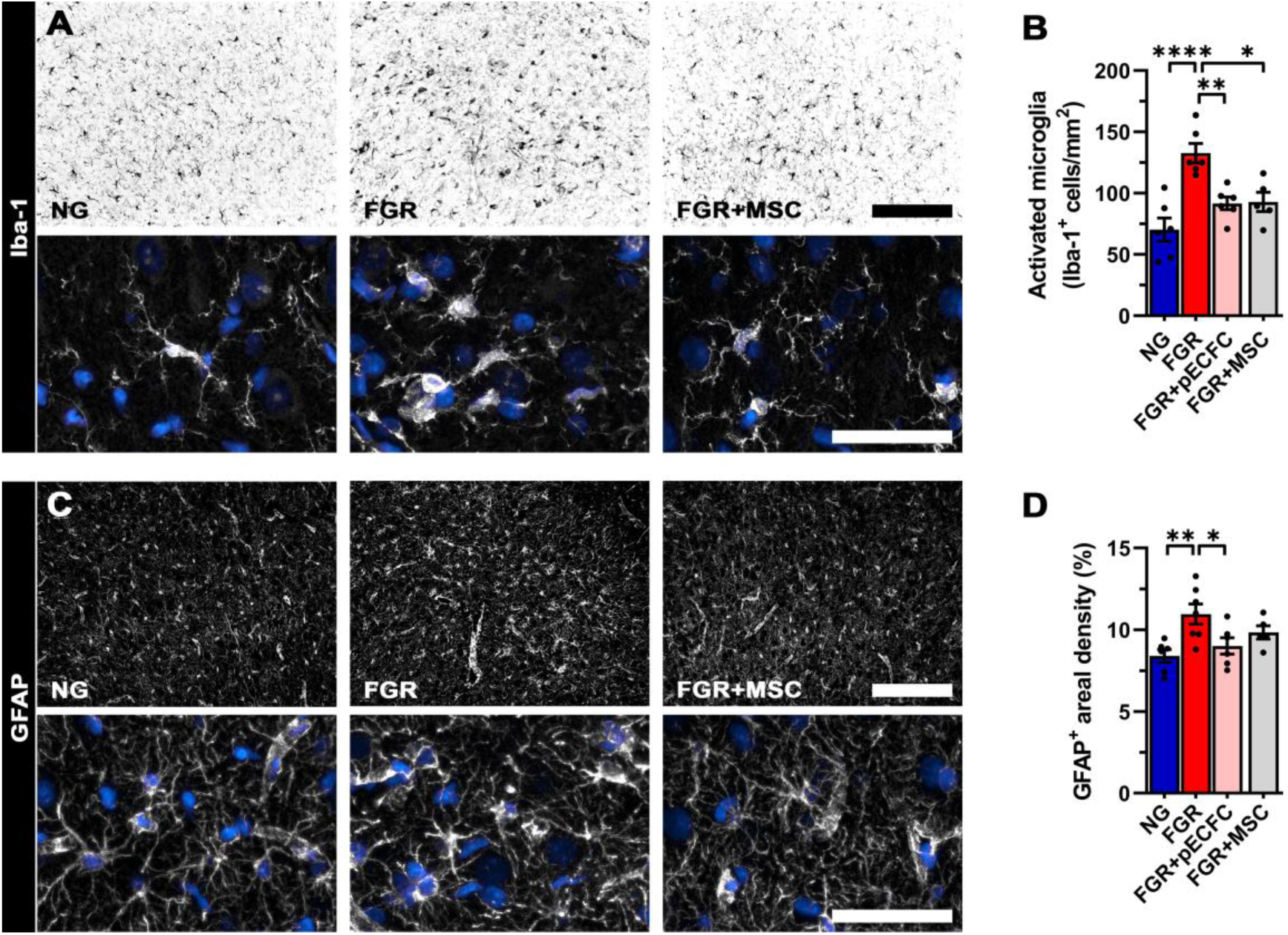
Mesenchymal stem cell treatment attenuates microglial but not astrocyte activation in FGR brains. **(A)** Representative staining of microglial expression (Iba-1; black) in the parietal cortex. **(B)** Treatment with MSCs significantly ameliorated the number of activated microglia. **(C)** Astrocytes (GFAP; white) showed evident alterations in morphology. FGR displayed activated astrocyte morphology compared with resting morphology of NG. **(D)** MSC treatment did not reduce astrocyte activation as assessed with aerial coverage of GFAP-positive labelled regions compared with untreated FGR. All values are expressed as mean +/- SEM (minimum n=5 for all groups). Two-way *ANOVA* with Tukey post-hoc test (*p<0.05, **p<0.01, ****p<0.0001) (Scale bars: 50 µm; low magnification: 200 µm).

**Supplementary Fig. 4.**
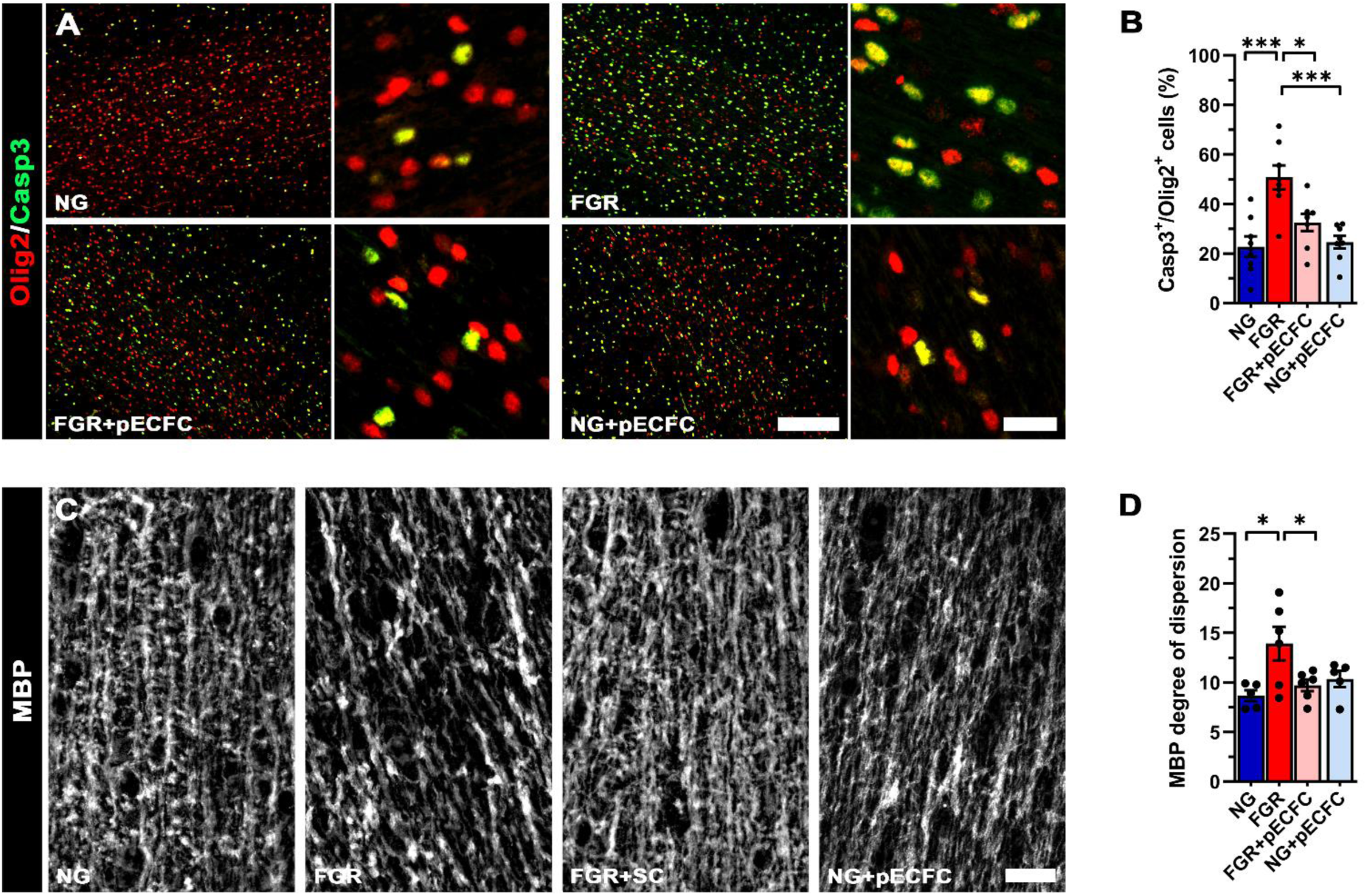
pECFC administration reduces oligodendrocyte apoptosis and improves white matter orientation. (**A**) Increased apoptosis (Casp3; green) of pan oligodendrocytes (Olig2; red) in the IGWM of FGR brain at postnatal day 4. (**B**) Administration of pECFCs significantly decreased apoptosis of oligodendrocytes in FGR compared with untreated FGR. (**C**) Representative labelling of MBP to demonstrate orientation of myelination along axonal fibers in the IGWM. FGR display more diffuse and disorganised labelling patterns compared with all other groups. (**D**) Directionality analysis showed an increase in dispersion of MBP orientation in the IGWM. FGR+pECFC displayed less variable dispersion comparable to that observed in NG and NG+pECFC groups. All values are expressed as mean +/- SEM (minimum *n*=5 for all groups). Two-way *ANOVA* with Tukey post-hoc test (*p<0.05, ***p<0.001) (Scale bars: 50 µm; low magnification A: 200 µm).

**Supplementary Fig. 5.**
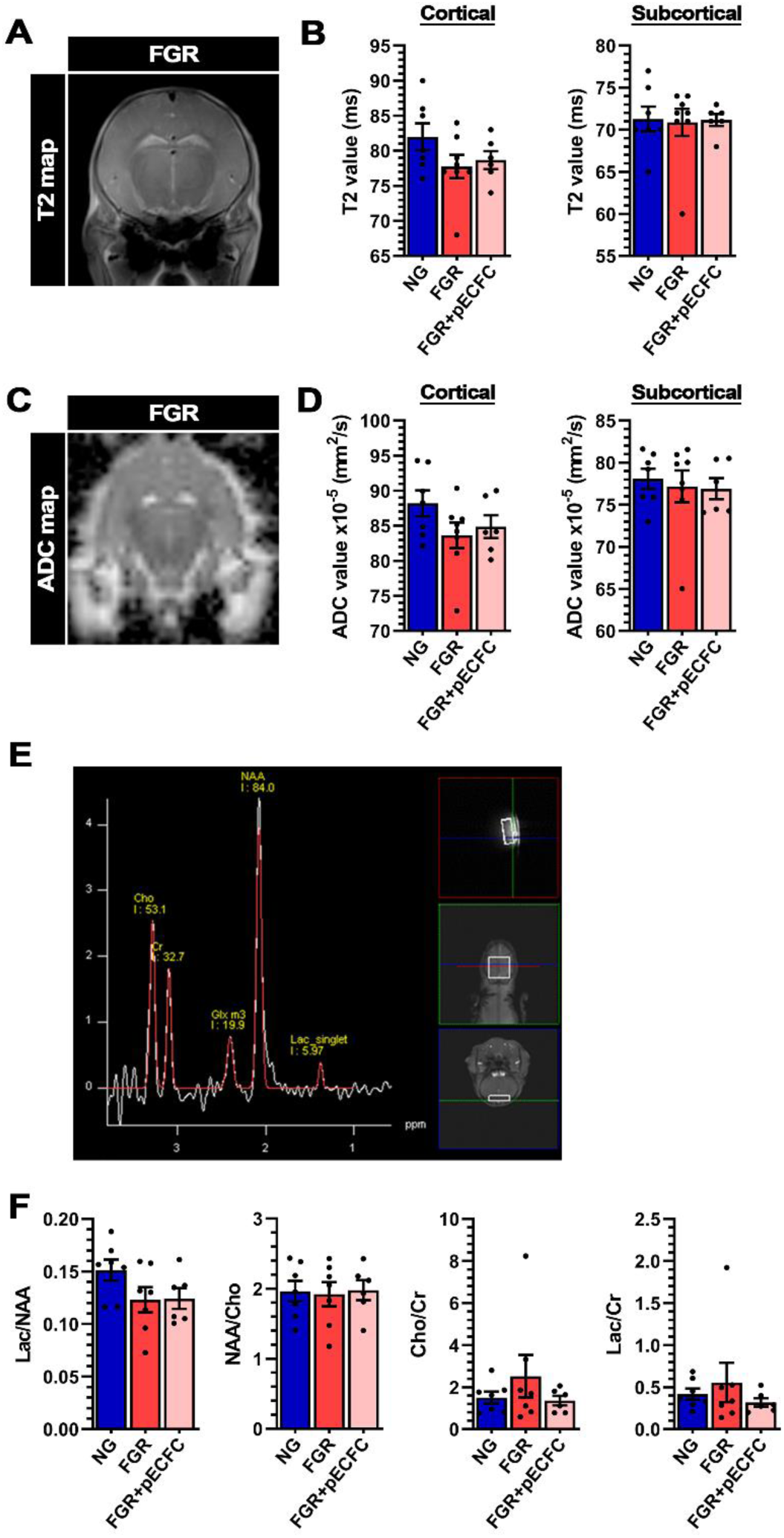
MRI and MRS analyses found no overt differences between NG, FGR and FGR+pECFC animals at day 4. Exemplar coronal views of MRI T2 maps. (***A***) and ADC maps (**C**) from an FGR brain at P4. Analysis of T2 and ADC acquisitions showed no significant differences between groups in the cortical or subcortical regions (***B*** *& **D***). (***E***) Representative snapshot of spectrogram showing the fronto-parietal brain voxel used to study brain metabolites. (*F*) No significant differences in metabolite ratios was observed between NG, FGR, and FGR+pECFC animals. Lactate (Lac), N-Acetylaspartate (NAA), Choline (Chol), Creatine (Cr).

## Notes

### Competing Interest Statement

The authors have declared no competing interest.

